# Quantifying uncertainty in brain-predicted age using scalar-on-image quantile regression

**DOI:** 10.1101/853341

**Authors:** Marco Palma, Shahin Tavakoli, Julia Brettschneider, Thomas E. Nichols, for the Alzheimer’s Disease Neuroimaging Initiative

## Abstract

Prediction of subject age from brain anatomical MRI has the potential to provide a sensitive summary of brain changes, indicative of different neurodegenerative diseases. However, existing studies typically neglect the uncertainty of these predictions. In this work we take into account this uncertainty by applying methods of functional data analysis. We propose a penalised functional quantile regression model of age on brain structure with cognitively normal (CN) subjects in the Alzheimer’s Disease Neuroimaging Initiative (ADNI), and use it to predict brain age in Mild Cognitive Impairment (MCI) and Alzheimer’s Disease (AD) subjects. Unlike the machine learning approaches available in the literature of brain age prediction, which provide only point predictions, the outcome of our model is a prediction interval for each subject.

## 1. Introduction

The process of brain ageing is known to be associated to a general decline in cognitive functions and higher risk of neurodegenerative diseases (Yankner et al., 2008; Denver and McClean, 2018). In some cases, both ageing and dementia affect the same areas in the brain (Lockhart and DeCarli, 2014). For these reasons, a deeper understanding of brain ageing in healthy conditions could potentially improve the diagnosis of neurodegeneration at early stages.

Neuroimaging provides a non-invasive and safe way to study brain structure and functioning. A large part of the research in neuroimaging data analysis has been focused on explanatory analyses aimed at describing the relationship between the brain and some variables of interest (such as neurodegenerative diseases, sex, physical activity). With the advent of large imaging databases, a prediction-oriented focus has been also considered, in order to detect individual differences among subjects that could be used in clinical practice (for example Yoo et al., 2018; Zhou et al., 2019).

The study of brain ageing has recently gained attention in the neuroscientific community thanks to the availability of this large amount of data and of computational tools for their analysis. A growing body of research employs neuroimaging to develop a biomarker of individual brain health, called “brain age” (Franke and Gaser, 2019; Cole et al., 2017). In the absence of a clear definition and assessment of biological brain age, a brain-derived prediction of chronological age is considered. In order to be integrated in clinical practice, a brain age biomarker should be easily accessible from brain data (or better, images), harmless for the subjects, computationally not demanding and correlated with other brain health indicators (Franke and Gaser, 2019). In addition, since there is a high variability between subjects in terms of their brain ageing, a useful biomarker should predict cognitive decline better than the chronological age itself.

In this work we propose a statistically grounded workflow that produces brain age individual predictions from 3-dimensional brain images. Furthermore, we go beyond simple point predictions by also providing prediction intervals of the brain age to quantify the uncertainty. Our model is trained on a control group with no ongoing brain diseases in order to avoid spurious effects due to other conditions. The same model can be used to predict age in neurodegenerative diseases, in order to provide a “baseline” or “normative” brain age, whose difference from the individual chronological age (brain-predicted age difference or *brainPAD* as in Cole et al., 2017) might inform about the extent of the effect induced by the pathology.

In addition, the prediction interval approach offers another potential binary biomarker (whether the chronological age falls within it). Since the width of the prediction interval is different for each subject, the same brainPAD could be interpreted in different ways in light of its location with respect to the individual prediction limits. The joint use of point and interval brain age predictions could therefore be employed to easily assess departures from a typical ageing profile.

The approach developed in this paper is based on modern statistical tools. In order to use 3D brain images without the need to summarise information by regions of interest, a functional data analysis (FDA) framework is adopted (Ramsay and Silverman, 2005; Horváth and Kokoszka, 2012). Functional data get this name because the observation for each statistical unit is a function^2^ (a curve, surface, or image). These data are usually considered as infinite dimensional and intrinsically continuous, even if the data collection process reduces them to a discrete series of observed points (Ramsay and Silverman, 2005, Section 3.2). In other words, the whole function is considered as the object of interest, and not only the specific value observed at a discrete location for each image. A common model in FDA is scalar-on-function regression (see Morris, 2015; Reiss et al., 2017 for reviews), which provides an effective way to predict a scalar quantity of interest from a functional observation, by fitting a regression model using the whole function as a covariate. In our context we call it *scalar-on-image regression*. The non-identifiability problem (Happ et al., 2018) arising from having sample size lower than the number of voxels for each image can be attenuated by imposing some assumptions on the data generating process (for example smoothness).

We obtain prediction intervals by integrating the FDA framework with quantile regression (Koenker and Bassett, 1978; Koenker and Hallock, 2001), a model that is largely used in fields such as economics (Fitzenberger et al., 2013) and ecology (Cade and Noon, 2003) to derive a more complete picture of the relationship between a covariate and the response variable. Quantile regression does not model the expected value (or a function of it) of the outcome of interest given the predictors, but some selected quantiles of the conditional distribution (for example the median). This model can be adapted for functional covariates: in a functional quantile regression model we explore the linear relationship between a certain quantile of the outcome and the 3D image. By fitting several quantile regression models we can build the prediction intervals given the covariates. Prediction intervals from quantile regression (or similar models) have received some attention in recent decades (Zhou and Portnoy, 1996; Meinshausen, 2006; Mayr et al., 2012), but not within the framework of functional data. In addition, the scalar-on-image quantile regression generates a regression coefficient with the same dimensionality as the brain image, providing an interpretable map that shows how the changes in each brain structure are related to the predicted age.

Our FDA-based approach departs considerably from other methods that are commonly used in the neuroimaging literature. The current state-of-the-art method in neuroimaging data analysis is the so-called *mass-univariate* approach implemented in the *Statistical Parametric Mapping* software (Ashburner et al., 2014). A model is fitted to predict the signal at each voxel independently using the clinical or demographic information as covariate, then a significance map is produced (see for further details Friston et al., 1994; Penny et al., 2011). Although computationally efficient, this approach does not explicitly model the spatial correlation of adjacent pixels and is not tailored for prediction purposes (Reiss and Ogden, 2010). The functional data approach allows instead the incorporation of the spatial structure by using smoothing techniques and in this way the fit of a global model for a scalar outcome given the entire brain image.

Another popular approach is based on machine learning algorithms. Franke and Gaser (2019) review a collection of studies published in the last decade based on a technique called relevance vector regression. They review a number of studies that examine associations with brain age, including effects of meditation and playing an instrument. Cole et al. (2019) collects a larger number of studies dealing with brain age prediction conducted from 2007 to 2018 with different imaging modalities and pathologies. Many of them adopt support vector regression (as the ones listed in Franke et al., 2012; Franke and Gaser, 2019 or Sone et al., 2019) or more recently Gaussian processes and convolutional neural networks (Cole et al., 2017; Cole, 2017; Varatharajah et al., 2018; Wang et al., 2019). A comparison between the predictive performances of these methods is difficult due to the use of different datasets and different age ranges, but according to Cole et al. (2019) the choice of the algorithm does not seem to play a fundamental role. However, these approaches provide only a point prediction with little knowledge of the internal procedure that returned it, and in particular deep learning methods are often criticised as “black boxes”. Our approach attempts to provide a better picture of the set of information on which brain age is based, introducing a straightforward quantification of uncertainty and at the same time producing a visual display of the regions that are most relevant for the prediction. In addition, the features of each step of the workflow proposed here can be evaluated, therefore improving the interpretability of the results. This last aspect is crucial in medical sciences and is particularly welcome for predictive modelling in neuroscience (Scheinost et al., 2019).

Another important distinction with the available literature on brain age prediction relates to the imaging techniques used. Although several models use functional imaging or multiple modalities, a large share of studies focused on structural magnetic resonance imaging (MRI), in particular T1-weighted images, usually segmentated into grey and white matter. Unprocessed MR images have also been employed with success (Cole et al., 2017). In this work we still remain in the family of structural imaging but we use tensor-based morphometry (TBM) images, that are obtained after a transformation of standard MRI images. TBM images give information about relative volumes of brain structures with respect to a common template; for this reason the images are all spatially registered. TBM quantifies volumetric differences in brain tissue for each voxel and is therefore specifically aimed at assessing the level of local cortical atrophy which might help to study brain degeneration for different diseases (Hua et al., 2008). To the best of our knowledge, this is the first study addressing brain age prediction from TBM images. The dataset used in this manuscript comes from the Alzheimer’s Disease Neuroimaging Initiative (ADNI, Mueller et al., 2005).

The work is structured as follows. Section 2 gives an overview of functional data analysis and quantile regression. Section 2.4 introduces the plan of the analysis and discusses details of the implementation. The main characteristics of the ADNI dataset are described in Section 3, while the results of the analysis are reported in Section 4 in terms of the predictions, their robustness with respect to the choices of the parameters in the model and their correlation with standard cognitive measures. Finally, Section 5 discusses the main findings, summarises the work and briefly introduces further research directions.

## 2. Materials and Methods

### 2.1. Functional data analysis

Functional data are realisations of a random function *X* ∈ *L*^2^(*T*), the space of square–integrable functions *f* : *T* → ℝ, for which

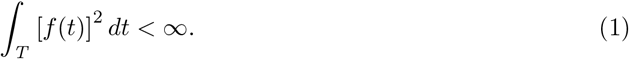

Typically in FDA we assume *T* ⊆ ℝ^*d*^ (Kokoszka and Reimherr, 2017; Ramsay and Silverman, 2005; Ferraty and Vieu, 2006). We define the inner product

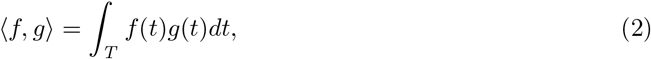

and the norm

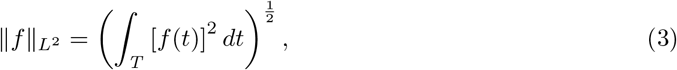

where *f, g* ∈ *L*^2^(*T*). The first order moment of *X* is the mean function 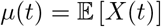; the second order variations of X are encoded in the covariance function

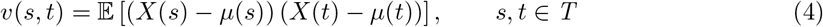

of which the variance function is a special case (*s* = *t*). A central object when dealing with functional data is the covariance operator, whose kernel is the covariance function *v*(*s, t*). It is defined as

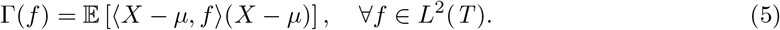

The covariance operator transforms a function *f* in another function Γ(*f*) whose values are

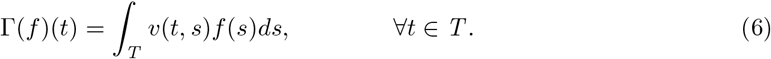

The covariance operator plays a key role in the Karhunen–Loève expansion for square–integrable functions,

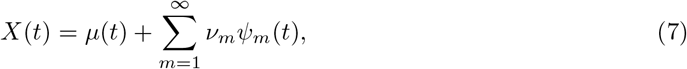

expressing *X* as an infinite linear combination of the deterministic eigenfunctions {*ψ_m_*(*t*)} of Γ with random and uncorrelated weights *ν*_*m*_. The eigenfunctions are the solutions of the eigendecomposition problem

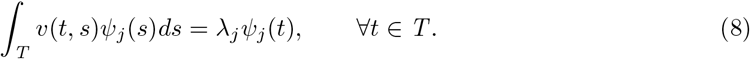

The eigenfunctions are orthogonal and rescaled to have unit norm, and their corresponding eigen-values {*λ*_*j*_} are in decreasing order.

The results of the eigendecomposition of the covariance operator can be interpreted under the framework of functional principal component analysis (FPCA), which aims at studying the principal modes of variation of the random function *X*. The eigenvalue *λ*_*m*_ is the part of the variance of *X* explained by the *m*-th eigenfunction, also called functional principal component. The random variables

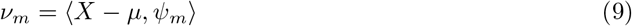

are called *scores*. The scores are uncorrelated and centered with variance *λ*_*m*_.

### 2.2. Quantile regression

Regression models are used to study the relationship between some fixed and known predictors *Z* = (*z*_1_, …, *z*_*M*_)^*T*^ ∈ ℝ^*M*^ and an outcome variable *Y*. For example, linear models are used to evaluate the change in the expected value of the continuous outcome conditioned on the values of the predictors, under specific assumptions on the error term. Nevertheless, there are occasions in which either these assumptions do not hold (for example, when there is heteroskedasticity in the residuals) or simply the main interest is to model specific quantiles of the conditional distribution of the response variable in order to produce a deeper analysis of the randomness of *Y* |*Z* that goes beyond the conditional mean^3^. Quantile regression (Koenker and Bassett, 1978) can effectively deal with these cases by specifying the model:

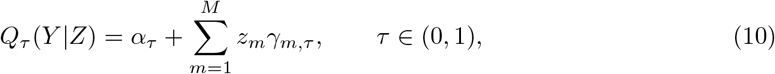

where *Q*_*τ*_ (*Y* |*Z*) is the *τ*-th conditional quantile of *Y* |*Z* defined as

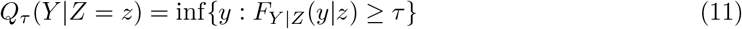

and

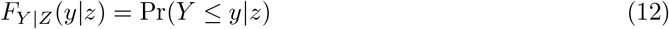

is the conditional cumulative distribution function of *Y* |*Z*. For example, *Q*_0.5_(*Y* |*Z*) is the median of the conditional distribution of *Y* |*Z*. The interpretation of *γ*_*m,τ*_ is similar to the one in linear models: it corresponds to the marginal effect on the conditional quantile due to a one-unit increment in the *m*-th covariate.

Given *n* observations, the estimation procedure for the model in Equation (10) is based on the following minimisation problem:

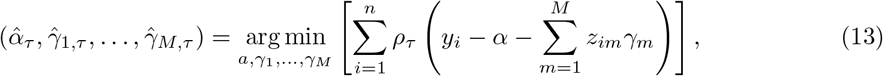

where 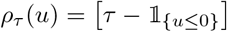 *u* is the check (or quantile loss) function (Koenker and Bassett, 1978). There is a relationship between the linear formulation *Y* = *Zγ* + *ε* and the quantile formulation in Equation (10). Under a linear data generating process *Y* = *α* + *Zγ* + *ε* with known *α* and *γ*, we can write the conditional quantile restriction

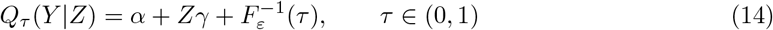

with *ε* being the mean zero random term of the model with cumulative distribution function (CDF) *F*_*ε*_. In this simple setting, the marginal effect of the covariate is constant across quantiles. Note that the result in Equation (14) holds for any distribution of the error term. Quantile regression can nonetheless accommodate more complicated data generating processes, like for example the location-scale model where *ε* is replaced by *σ*(*Z*)*ε*, with *σ*(*Z*) > 0 and *ε* ⊥ *Z*. In this case the variance of the random term depends on Z and it can be shown that the estimated slope in the quantile regression model will be governed by the quantiles of *ε*.

All the quantile regression models return as output a prediction at a specific quantile level. For example, the model with *τ* = 0.5 gives the conditional median prediction for each experimental unit given particular values of the covariates. Predictive accuracy of the conditional median can be measured through the mean absolute error (MAE) and the root mean square error (RMSE) between the point predictions and the observed responses. By fitting a model for several values of *τ*, we can also build prediction intervals for new observations (*y*\*, *z*\*) (Davino et al., 2013; Mayr et al., 2012). For example, if we fit a model on the same data for two quantile levels *τ*_1_ = *δ/*2 and *τ*_2_ = 1 − *δ/*2 (with *δ* ∈ (0, 1)), the interval

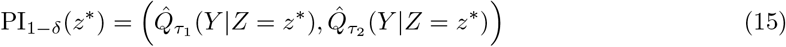

should contain the observed response value for new data (1 − *δ*)100% of the time (provided Equation (10) is true). For example, a 90% prediction interval can be obtained by fitting a model for *τ*_1_ = 0.05 and *τ*_2_ = 0.95. This prediction model can effectively handle heteroskedasticity or skewness, since in quantile regression there are no assumptions on the response distribution: using simulated data Davino et al. (2013) provide examples in which prediction intervals obtained via quantile regression achieve the nominal levels where ordinary least squares prediction intervals fail. This is also confirmed theoretically in Zhou and Portnoy (1996): the coverage probability tends to 1 − *δ* with an error of *O*(*n*^−1*/*2^), as the sample size of the training set *n* → ∞.

### 2.3. Functional quantile regression

A large body of literature has been developed in order to translate regression models into the functional framework. For example, functional GLMs are now well established in the theory, both in the frequentist and Bayesian approaches (Müller and Stadtmüller, 2005; Crainiceanu et al., 2009). Quantile regression (Koenker and Bassett, 1978) has also been extended in the functional data paradigm: first with Cardot et al. (2005), then with Kato (2012) and Yao et al. (2017), the model has been readapted for the case of functional covariates with scalar response. The model illustrated in Kato (2012) shares the main characteristics with the scalar-on-function regression of Müller and Stadtmüller (2005), except for the assumption that the conditional quantile is a linear function of the (centered) covariates. In particular, the conditional quantile of the response is expressed as a linear function of the scalar product between the functional data and a coefficient function *β*_*τ*_ (·) ∈ *L*^2^(*T*):

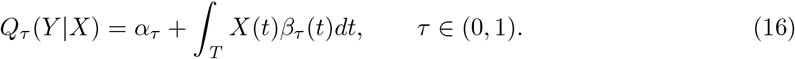

The functional nature of the coefficient makes its interpretation less straightforward than in standard regression. In the regions where *β*_*τ*_ (*t*) = 0 any increment in the covariate produces no marginal change on the quantile of the conditional distribution *Y* |*X*. On the other hand, if *β*_*τ*_ (*t*) is constant over a region *T* * ⊂ *T* and null elsewhere, then only the region *T* * plays a role in the prediction of the conditional quantile. Despite the differences between quantile and linear scalar-on-function regression, the same difficulties of the interpretation of the functional coefficients discussed in James et al. (2009) apply. The model can easily accommodate scalar covariates *z*_1_, …, *z*_*P*_ (see for example Yao et al., 2017):

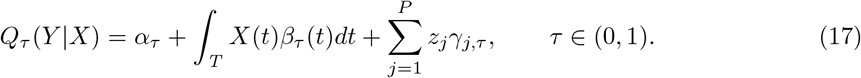

In order to estimate the parameters in Equation (16), both the predictors and the coefficient functions are represented in the truncated Karhunen–Loève expansion in Equation (7):

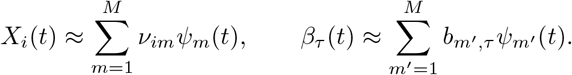

Thanks to the orthonormality of the eigenfunctions *ψ*_*m*_,

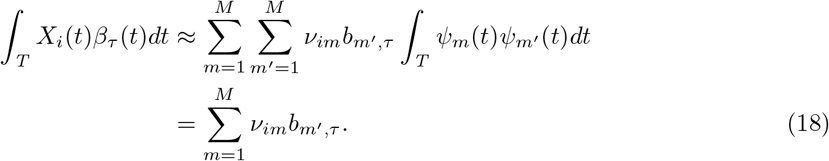

Thus the functional model in (16) becomes a standard quantile regression problem of the form

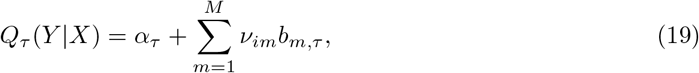

where *α*_*τ*_ and *b*_1,*τ*_,…, *b*_*m,τ*_ are estimated as in Equation (13). The estimated functional coefficient is then reconstructed by computing

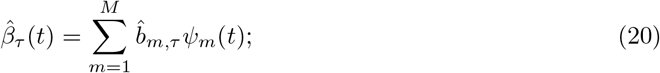

for a given *τ* the estimated value for the quantile function is obtained by plugging in the estimated coefficient into (16):

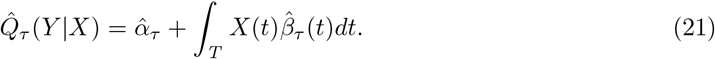

In this functional principal components regression (FPCR) setting, the number of principal components *M* to be used as regressors controls the smoothness and the approximation error with respect to the real images. The choice of *M* could be automated by using information criteria or percentage of variance explained; nevertheless, there is no guarantee that the first *M* components (which explain the most of the variability of X) are also able to capture effectively the relationship between the functional predictor and the scalar response (Febrero-Bande et al., 2017; Delaigle and Hall, 2012). For this reason, a simple option could be to select *M* such that a very large share of explained variability is represented and then use LASSO regularisation within the quantile regression model (Belloni and Chernozhukov, 2011; Wang, 2013). The regularisation might produce a different subset of selected variables across different quantile levels *τ*. Since for each *τ* a different model has to be fitted, the plug-in estimator 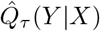 is not guaranteed to be monotonically increasing in *τ* as the conditional quantile function 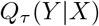 is by construction.

It must be considered that the bias introduced by the penalised estimation could harm the interpretability of the coefficients for each covariate. A way to solve this issue is the post-*l*_1_ quantile regression, where LASSO is used only for model selection and then a vanilla quantile regression model is fitted using only the covariates selected. This approach guarantees better convergence rates and could reduce the bias (Belloni and Chernozhukov, 2011).

### 2.4. Data analysis workflow

#### 2.4.1. Imaging

The brain images are acquired using structural MRI. This workflow does not depend on any specific preprocessing stages, except for intersubject registration to an atlas image, such that voxels from different images are aligned.

More transformations can be operated on the structural MR images. For example, the analysis can be based on tensor-based morphometry (TBM) images. TBM is an image technique that aims at showing local differences in brain volume from structural imaging. In a cross-sectional setting (one image for each subject), each image is aligned to a common MRI template called *minimal deformation template* (MDT). The deformation induced by this alignment can be represented by a function that maps a 3-dimensional point in the template to the corresponding one in the individual image. The Jacobian matrix of the deformation can be used to inform about volume differences in terms of shearing, stretching and rotation. The determinant of the Jacobian matrix for each voxel is then a summary of local relative volumes compared to the MDT: a value greater than 1 indicates expansion, while a value less than 1 means contraction. Further details about TBM are available in Ashburner and Friston (2004).

In order to reduce the dimensionality of the problem, the voxels outside the brain can be excluded from the analysis imposing a mask on the images. We used FSL (through its R interface `fslr`, Muschelli et al., 2015) to obtain a mask on the template image with smooth boundaries.

#### 2.4.2. Basis expansion

A common assumption in FDA is that the observed data are a noisy, discretised version of the true underlying signal function that is of interest in the analysis. In other words, the values observed at a specific voxel may be contaminated with some measurement error that could have an impact on the spatial correlation structure within the images. Removing this measurement error leads therefore then to smoother images, improving the performances of FPCA.

For this reason, nonparametric basis expansion techniques such as B-splines or wavelets are usually employed. The latter are chosen mainly when the underlying function is thought to be characterised by rapid changes in behavior (Ramsay and Silverman, 2005); B-splines are instead preferred for their properties (compact support, unit sum) when less abrupt changes in the function are expected. In this case, TBM images are already smooth by construction, so we can use B-spline basis functions with the main aim to obtain a parsimonious representation (under the fairly safe assumption that the main sources of error have been already removed).

In order to get a 3-dimensional basis function, a tensor product of univariate B-spline basis functions is considered. Denote by 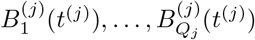 the univariate basis functions for the *j*-th dimension (*j* = 1, 2, 3). The number of basis functions for each dimension is *Q*_*j*_ = *l*_*j*_ + *r* − 1, where *l*_*j*_ is the number of knots and *r* is the degree of the spline. We now define the set of basis functions

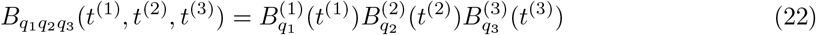

for *q*_*j*_ = 1,…, *Q*_*j*_, for *j* = 1, 2, 3.

In order to derive the projection of each image onto this set of basis functions, we define the following matrix of basis functions using the Kronecker product

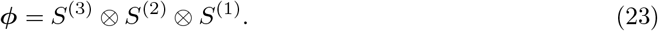

where *S*^(*j*)^ is the *P*_*j*_ × *Q*_*j*_-dimensional matrix whose *q*_*j*_-th column contains the evaluation of the function 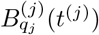 at each point *t*^(*j*)^ (for *j* = 1, 2, 3) and *P*_*j*_ is the number of points for the *j*-th dimension. The matrix *φ* has dimensions *P*_1_*P*_2_*P*_3_ × *Q*_1_*Q*_2_*Q*_3_ (the number of rows is equal to the number of voxels and the number of columns is equal to the number of basis functions). Once the basis set is determined, this can be used as set of regressors where the original (vectorised) image is the response variable. Estimation can be performed via ordinary least squares:

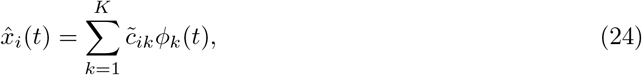

where *K* = *Q*_1_*Q*_2_*Q*_3_, 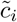 is the *K*-dimensional vector containing the coefficients of the projection for the *i*-th image and *φ*_*k*_(*t*) is the *k*-th basis function. In compact form, all the *N* images are represented by the product of the *N* × *K* coefficient matrix 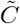 and the matrix of basis functions *φ*. We center the projected data (equivalent to centering the raw data since the projection is linear). This apparently negligible aspect is actually very relevant in the big data context as it allows to parallelise the basis expansion stages without the need to import and store simultaneously all the images. We call the centered coefficient matrix *C*.

In this work we used a 3D tensor product of quadratic B-spline univariate basis functions with equidistant knots. The number of knots (or analogously their spacing) can be fixed in advance, but a poor choice might heavily affect the number of basis functions that are needed to represent the functions and consecutively the computational time and the quality of projection. For this reason a preliminary study on a subset of the data is recommended. Outcomes of interest for this preliminary study could be the number of non-zero basis functions within the masked image, the average time needed for the projection of an image and the *R*^2^ value obtained from the regression of each image using as design matrix the matrix of basis functions. The latter value can be interpreted as a proportion of variance explained. At this stage, it is highly recommended to retain as much variability as possible: a 0.95 threshold for *R*^2^ should work for many applications and should ensure a manageable set of basis functions. Alternative criteria could be established in terms of full width at half maximum (FWHM).

#### 2.4.3. Functional PCA

The coefficients of the projection are the quantities needed to solve the eigendecomposition problem in Equation (8). In this section, we rely heavily on Ramsay and Silverman (2005, Section 8.4.2), with minor modifications to make this high dimensional problem computationally feasible. The procedure is described also in Chen et al. (2018).

The sample variance-covariance function can be written as

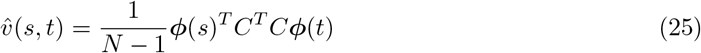

using the same decomposition in (24). Suppose then that the eigenfunctions in Equation (8) can be expressed as linear combinations of the same basis functions *φ*:

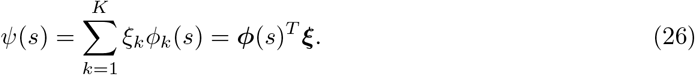

Then the eigenanalysis of the covariance operator described in Equation (8) takes the following form:

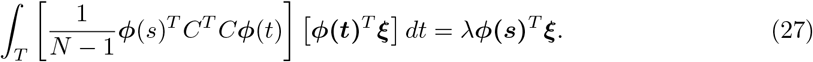

Denoting by *W* the *K* × *K* symmetric basis product matrix with elements

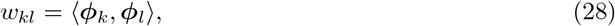

Equation (27) can be rewritten as

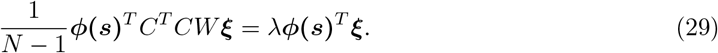

The entries in *W* are usually computed with some numerical quadrature rules (Ramsay and Silverman, 2005) but these procedures are computationally demanding in our 3D context. The cross product, although less accurate at the boundaries with respect to the trapezoidal rule, offers a good result in shorter time. Simplifying both sides of Equation (29) by *φ*(*s*)^*T*^ (the relationship must hold for all *s*) we obtain

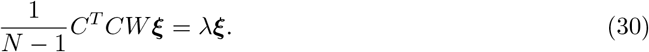

In order to get orthonormal eigenfunctions, some constraints must be imposed:

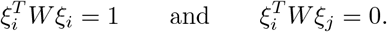

These are fulfilled by setting *u* = *L*^*T*^ *ξ*, where *L* is obtained through the Cholesky decomposition *W* = *LL*^*T*^ (Ramsay and Silverman, 2005, p. 181); solving the equivalent problem

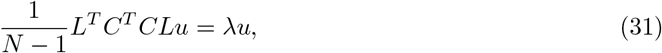

the original eigenfunctions are obtained using *ξ* = (*L*^*T*^)^−1^*u*.

We note that for *A* = (*N* − 1)^−1/2^*CL* the eigendecomposition problem consists in finding the eigenvalues and eigenvectors of *A*^*T*^ *A*. These can be obtained in a computational efficient way by using the SVD of the matrix *A*. In particular, the non-zero eigenvalues *λ* are equal to the squared non-zero singular values, whereas the eigenvalues *u* of *A*^*T*^ *A* are equal to the right singular vectors of *A*. The *m*-th score for the *i*-th image is then

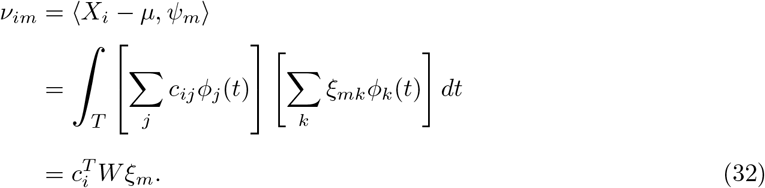

#### 2.4.4. Functional Quantile Regression

The scores obtained after FPCA are plugged into a standard quantile regression problem. We create the design matrix for the quantile regression model using the first *M* scores for each image such that the first *M* eigenfunctions represent at least 80% of the variability within the sample (see Section 4.3 for a sensitivity analysis). LASSO regularisation can be applied within the quantile regression framework. The minimisation problem in Equation (13) can be readapted therefore to our situation by writing

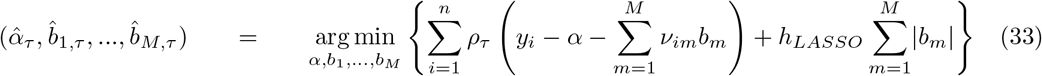

where *h*_*LASSO*_ is the LASSO tuning parameter. For a specific value of *h*_*LASSO*_, a solution path is found, where the Lasso penalty will induce the shrinkage of the estimates towards zero, but also sparsity, as some estimates are exactly zero (Tibshirani, 1996).

Several R packages offer built-in functions that perform automatic selection of the tuning parameter. For this purpose, we use the package rqPen (Sherwood and Maidman, 2017), that produces penalized quantile regression models for a range of tuning parameters and then selects the one with minimum cross-validation error.

#### 2.4.5. FPCA and functional quantile regression in a prediction setting

The scores are obtained by taking an inner product of each image with the eigenfunctions estimated on the training set. For this reason, they can be obtained for images from other datasets with the same formula, even if the properties of zero mean and variance equal to the eigenvalues apply only for the training dataset. The scores are in turn produced within the FPCA step, where the estimation of the eigenfunctions depends on the training data as well.

This workflow is aimed at deriving brain age prediction intervals for healthy individuals. This means that FPCA and functional quantile regression should be based on a dataset of control subjects. In order to get predictions for this dataset, 10-fold cross validation can be used, reducing in this way the risk of overfitting. Age predictions for subjects with neurodegenerative diseases can be obtained from the same normative model. In this case the full dataset of control subjects can be used for FPCA and functional quantile regression and the brain age is to be interpreted as the equivalent brain age of a healthy individual having the same brain image.

The R code implementing the workflow is available at https://github.com/marcopalma3/neurofundata.

#### 2.4.6. Alternative models

The degree of smoothing in the basis expansion step can be controlled in different ways, by changing either the location or the numbers of knots. When the number of knots is equal to the number of voxels, we recover the original data, where the coefficient of the basis functions are just the observed values at each voxel. The analysis of the “unsmoothed” images can still be based on standard multivariate analysis techniques such as PCA and quantile regression, but it requires an increased computational effort. The data matrix containing the images as rows is indeed large (in our case the memory needed to store it is more than 6.4GB) and high performance computing tools are required to fit models on these data. In addition, quantile regression under memory constraints is receiving attention only recently (Chen et al., 2019), therefore the calculation of the prediction interval is not straightforward. A small amount of smoothing is recommended to reduce both the storage issues and the computational time required to train the model.

## 3. Data

The workflow proposed in Section 2.4 is applied on a dataset coming from the Alzheimer’s Disease Neuroimaging Initiative (ADNI, Mueller et al., 2005), that supports the investigation about biological markers to be used to detect Alzheimer’s Disease (AD) at early stages. The sample used in this paper is made of 796 subjects, identified through an ID code, for which several demographic and clinical variables are measured. In this analysis, we will consider only the chronological age at the entry of the study (ranging from 59.90 to 89.60 years; mean age 75.60±6.29) and their diagnosis: 180 subjects were diagnosed with AD, 387 with MCI (Mild Cognitive Impairment, considered as an intermediate stage between healthy condition and AD) and 229 people were belonging to a control group of cognitively normal (CN) subjects. The histogram of age by diagnosis group is displayed in Figure 2.

**Figure 1:**
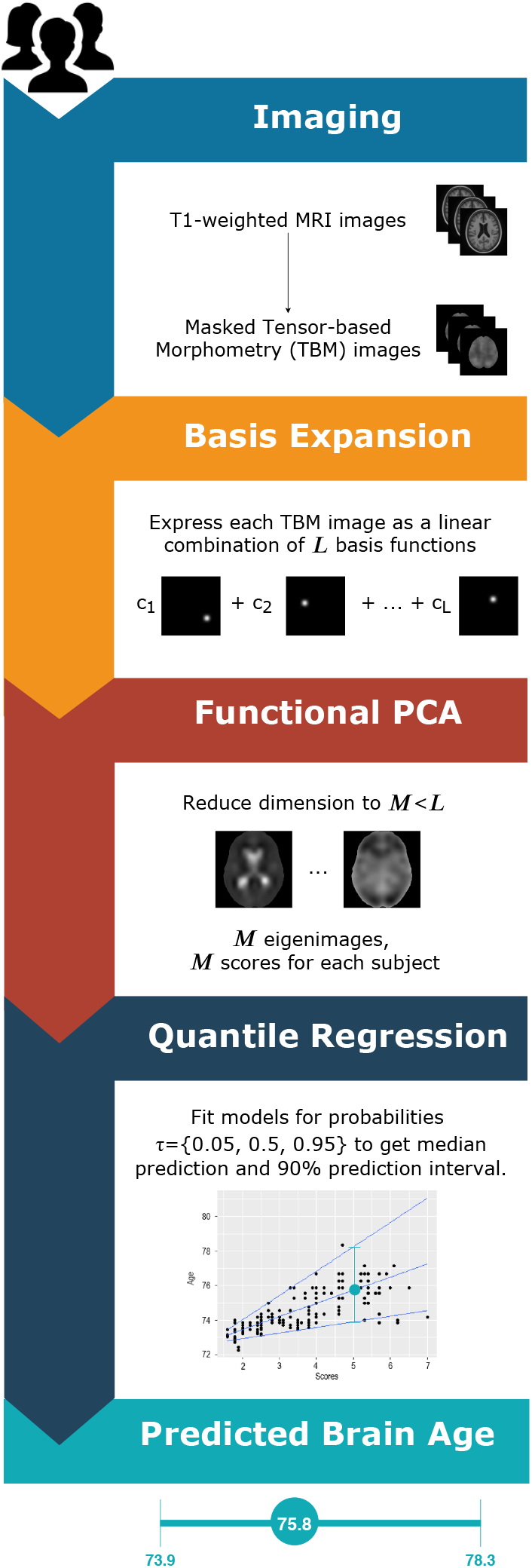
Flowchart of the analysis from the brain images to the predicted intervals.

**Figure 2:**
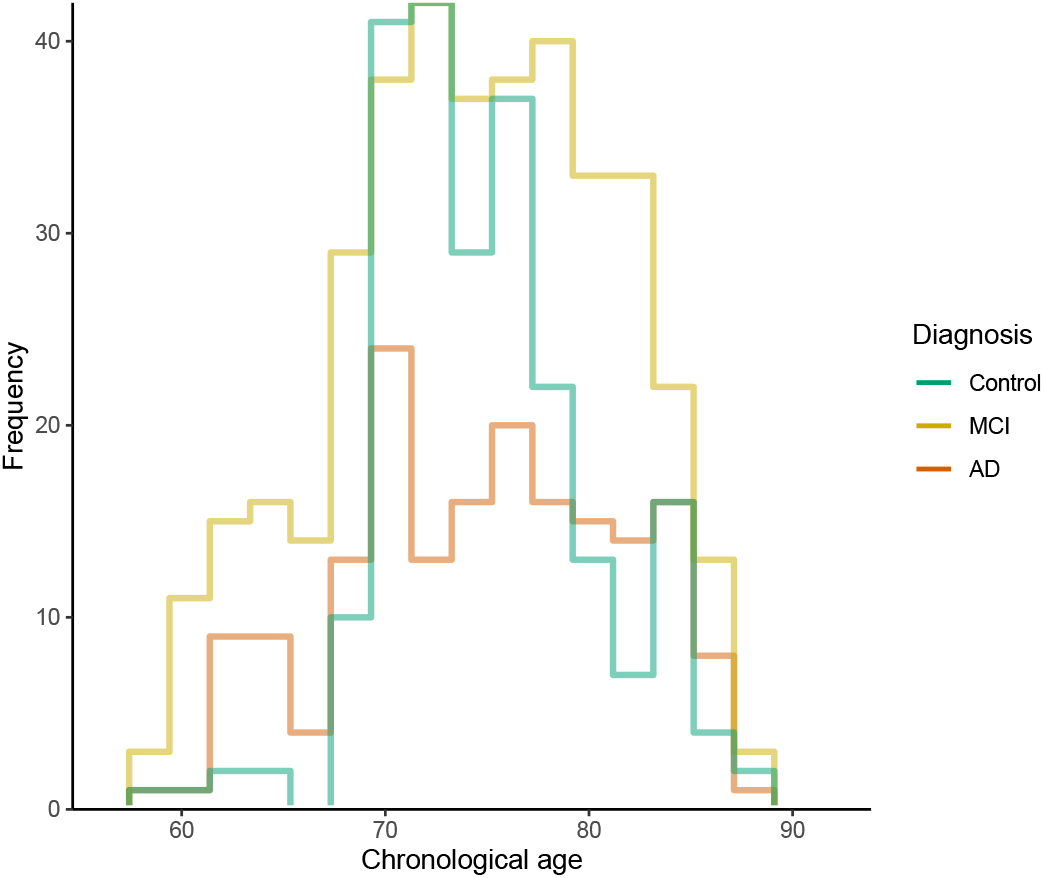
Histogram of age of the subjects in the sample, for each diagnosis. The number of bins has been fixed using the Freedman-Diaconis rule (Freedman and Diaconis, 1981).

The functional part of the dataset consists of tensor-based morphometry (TBM) images taken at the baseline of the study for each subject. In this dataset, the threshold 1 is rescaled to 1000 for computer number format reasons. Information about the preprocessing stages for the ADNI TBM dataset is available in Hua et al. (2013).

The analysis is based on the original 3D TBM scans (220 × 220 × 220, with voxel size equal to 1 mm^3^). The conventional neurological orientation (“right is right”) is used: the (*x, y*) axes of the images are set such that *x* increases from left to right and *y* increases from posterior to anterior.

The mean functions for each diagnosis are shown in Figure 3. MCI and AD patients share similar average brain volumes patterns (namely, expansion of the lateral ventricles and shrinkage almost everywhere else) even if the intensity of the expansion is higher for people with dementia. The expansion of the lateral ventricles is also visible in the healthy control mean function, but it is less pronounced. Conversely, the healthy control mean function shows other slightly expanded brain areas, such that the cerebellum and several regions in the posterior and frontal lobes. Further analyses based on the voxelwise variance functions per each group show that the lateral ventricles are the areas with the highest variability in terms of volume expansion.

**Figure 3:**
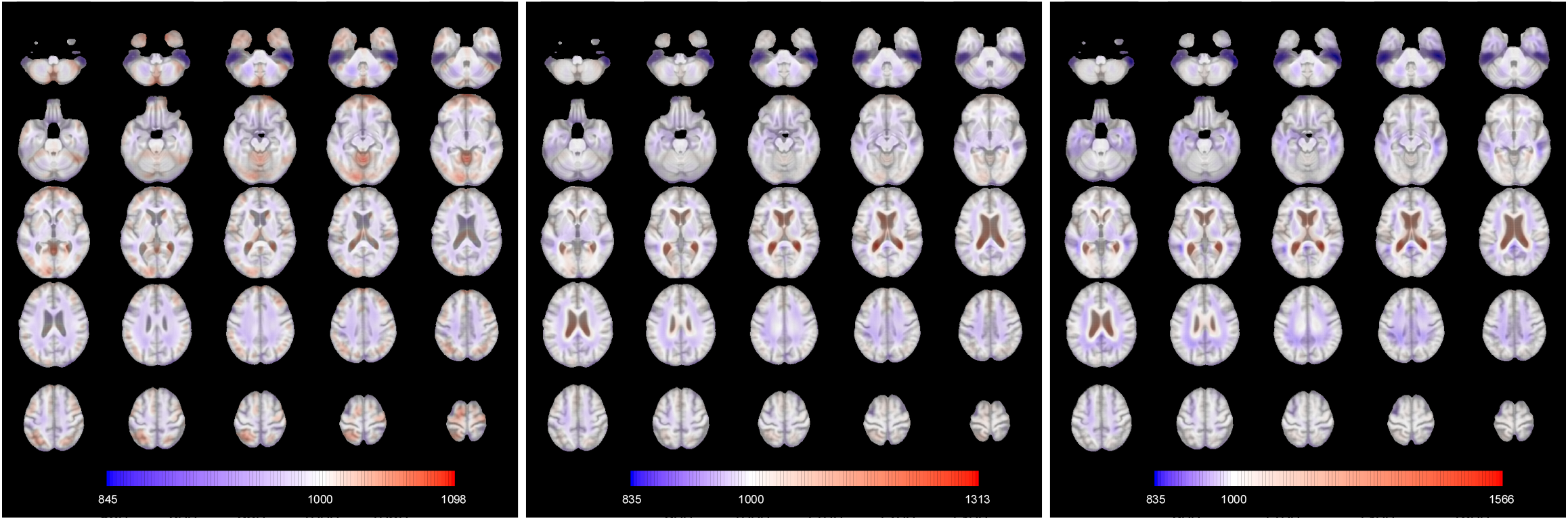
Axial slices of the mean images for each diagnosis (from left to right: Control, MCI, AD). Slices are ordered from bottom to top. The colours are overlaid on the corresponding slice of the MDT.

## 4. Results

### 4.1. Prediction accuracy

The preprocessed images are masked to remove unnecessary voxels for the analysis. A 3D smooth mask is obtained by smoothing the raw mask with a Gaussian kernel with standard deviation equal to 2 voxels (FWHM 4.7 mm) and thresholding it at 0.5, to regularise the boundary, producing just over 2 million nonzero voxels.

For the dataset at hand the B-splines projection with equidistant knots every 12 mm (equivalent to FWHM ≈ 15.33 mm) for each dimension allows to represent each image with *R*^2^ approximately equal to 0.96. The number of B-spline functions in the tensor product that fall within the mask is 2694. In the current implementation, the process of importing one image into R and obtaining its B-spline coefficients takes approximately 30 seconds.

The eigendecomposition problem in Equation (8) solved for the dataset of healthy control subjects returns M = 54 eigenfunctions of which the first 3 are plotted in Figure 4. In analogy with standard PCA, a basic interpretation can be provided. The first eigenfunction clearly distinguishes the lateral ventricles from the rest of the brain. Subjects with high scores for this eigenfunctions will show stronger expansion within the lateral ventricles with respect to the mean function. Due to the similarities with the observed patterns in the mean function for the subjects with disease, it is likely that the scores for this eigenfunction computed for all the 796 subjects in the dataset are correlated with the diagnosis and with the chronological age, for the known interplay of the effects of these two factors. The second mode of variation refers instead to a more general expansion across the whole brain: in other words, it discriminates between individuals with bigger brains and those with smaller ones. For this reason, this component might account for some sex-related effects, as males have on average larger overall absolute brain than females (Ruigrok et al., 2014). The third eigenfunction weights negatively some of the internal parts of the brain. This component might therefore roughly distinguish white matter from the cortex, even if this interpretation is not very clear and can be influenced by the smoothing induced by the projection onto the basis functions. The first 3 components account for 36.25% of the variance of the images of the healthy control group.

**Figure 4:**
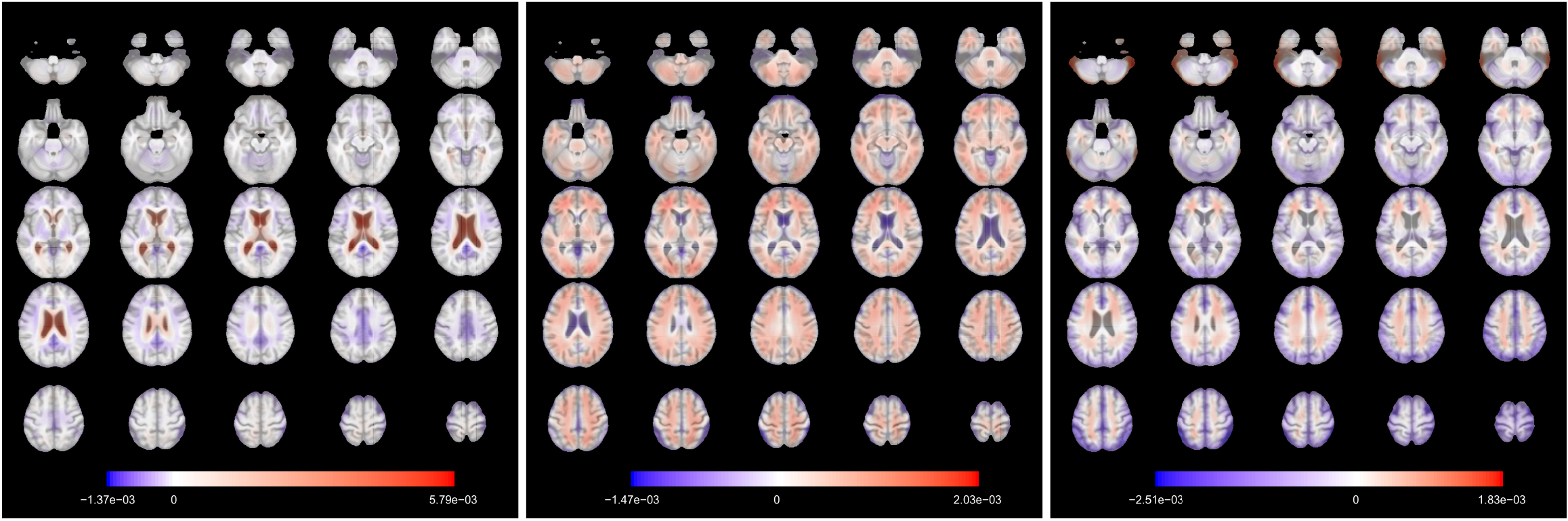
Axial slices of the first 3 eigenfunctions for the control subset. Slices are ordered from bottom to top. The colours are overlaid on the corresponding slice of the MDT. The eigenfunctions account respectively for 15.43%, 13.95%, 6.87% of the total variability. The signs of the eigenfunctions are determined on the basis of clinical interpretation.

We compute the scores for MCI and AD individuals as the product of the centered images and the eigenfunctions in Figure 4. For the control subjects, we use 10-fold cross validation (with check function as loss function) to run FPCA, produce scores and fit the models such the predictions are obtained on held-out data. Quantile regression models for *τ* ∈ {0.05, 0.5, 0.95} are considered. Table 2 shows that the MAE and RMSE based on the difference between median brain-predicted age and chronological age are lower for control subjects than the other groups. This result is expected under the choice of a normative model that predicts brain age in absence of any diseases and indicates that the two subpopulations (controls vs. cases) show different ageing characteristics (if they were belonging to the same population, the MAE and RMSE would have been similar).

**Table 1:** Summary statistics for each diagnosis group. *N* is the number of subjects in each group. The second part of the table shows selected quantiles of age.

| Diagnosis | $N$ | Min. | 1st Qu. | Median | Mean | 3rd Qu. | Max. |
| --- | --- | --- | --- | --- | --- | --- | --- |
| Control | 229 | 59.90 | 72.30 | 75.60 | 75.87 | 78.50 | 89.60 |
| MCI | 387 | 60.10 | 70.85 | 75.60 | 75.30 | 80.40 | 89.30 |
| AD | 180 | 59.90 | 70.98 | 76.15 | 75.90 | 81.58 | 89.10 |

**Table 2:** Summary of the prediction results by diagnosis. Cor: correlation between predicted brain age and chronological age. CI_*Cor*_: confidence interval for the correlation between predicted brain age and chronological age, obtained via Fisher-z transformation (Myers et al., 2013, Section 19.2). 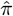: sample coverage (proportion of cases for which the 90% prediction interval contain the chronological age). *-pos: proportion of cases for which the chronological age is less than the lower limit of the 90% prediction interval.

| Diagnosis | $N$ | MAE | RMSE | Cor | 95% $CI_{Cor}$ | $\hat{\pi}$ | *-pos |
| --- | --- | --- | --- | --- | --- | --- | --- |
| Control | 229 | 3.49 | 4.43 | 0.48 | [0.37, 0.57] | 0.86 | 0.05 |
| MCI | 387 | 4.99 | 6.12 | 0.46 | [0.38, 0.54] | 0.68 | 0.24 |
| AD | 180 | 5.16 | 6.27 | 0.38 | [0.25, 0.50] | 0.64 | 0.28 |

The MAE observed for the control group is 3.49, in line with other results obtained in the literature for other MRI datasets and different age ranges (Cole et al., 2019). In addition, as shown in Figure 5, the smoothed regression line for control subjects indicates that the average *brainPAD* (difference between predicted and chronological age) is close to zero for the whole age range, while it departs from it for the other groups in the predicted age range between 73 and 75. Prediction metrics do not improve after debiasing using post-*l*_1_ quantile regression.

**Figure 5:**
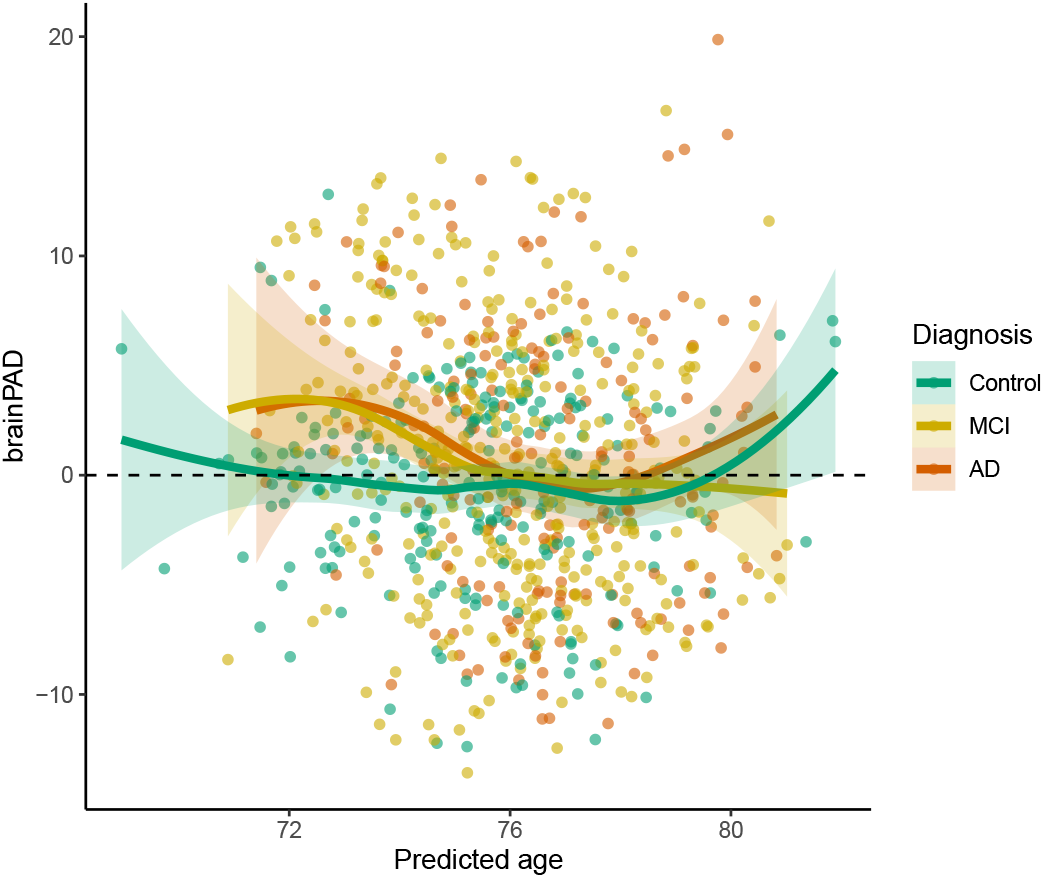
Plot of the brainPAD vs. predicted response. The coloured lines are local regression lines obtained with loess (locally estimated scatterplot smoothing) with span = 0.75 and 95% confidence bands.

We focus now our attention on the features of the 90% prediction intervals and the sample coverage. We observe that the actual sample coverage for control subjects is slightly lower than the nominal level. The groups with cognitive impairment show lower coverage with respect to the control group: the chronological ages of around 1 in 3 subjects with diseases do not fall in the prediction intervals obtained under the normative model. When we further analyse the direction of the discrepancy, we can define a “*-positive brainPAD” group (for which the chronological age is lower than the lower limit of the prediction interval, or equivalently with positive brainPAD and chronological age outside the prediction interval) and a “*-negative brainPAD” one (composed of those subjects with negative brainPAD and chronological age outside the prediction interval). While the share of *-negative subjects is approximately constant across the diagnosis, the percentage of *-positive subjects for MCI and AD groups is approximately 5 times the one for the control subjects. This result aligns with the literature, where it has been shown that MCI and AD patients show higher apparent brain age (Cole et al., 2019; Franke et al., 2012): for this reason the *-positive group is more interesting for their potential correlation with other disease indicators. All the prediction intervals are plotted in Figure 6, stratified by diagnosis and sorted by predicted age. The prediction intervals for the control subjects are scattered closer to the line of identity between predicted and chronological age and there are no relevant trends in the residuals that are left unexplained by the regression models. The variability of the width of the 90% prediction intervals is displayed in Figure 7: the average width is similar for the 3 diagnosis groups, but there is higher variability in the width distribution of the MCI and AD subjects. Moreover, *-positive brainPAD is mainly observed in the lower part of the age domain covered in the dataset. This could be just a consequence of our regression approach, or it might be due to the low number of subjects in the training set with chronological age less than 70, which might produce issues in the estimation of extreme quantiles of the conditional distribution of the outcome.

**Figure 6:**
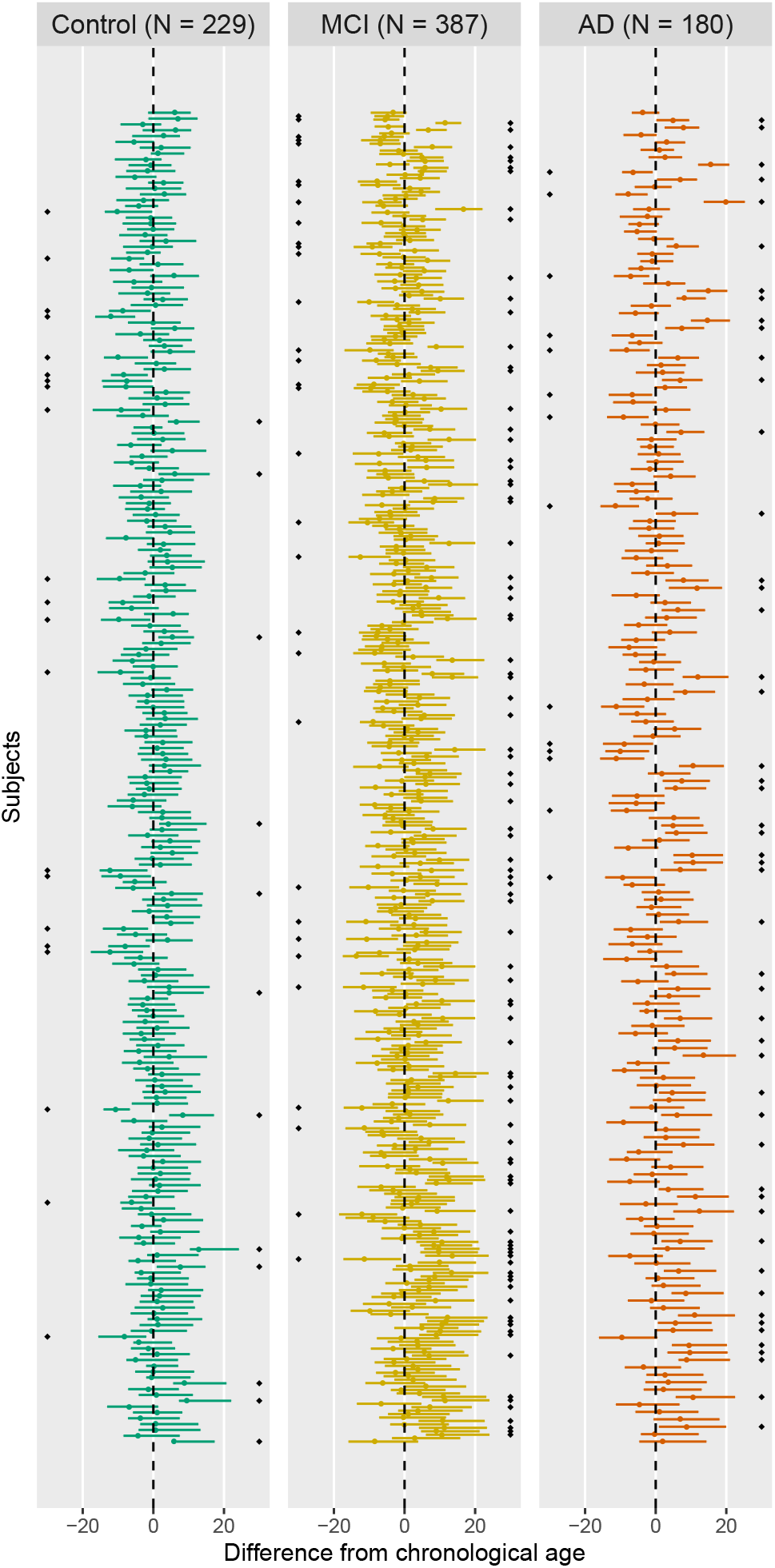
Brain age 90% prediction intervals, relative to chronological age. There is one interval per subject, and subjects are sorted in descending order of predicted brain age (higher predicted ages at top). The black diamonds indicate the subjects for which chronological age does not fall into the prediction interval; the side indicates if the subject is in the *-negative (diamonds on the left) or *-positive group (diamonds on the right).

**Figure 7:**
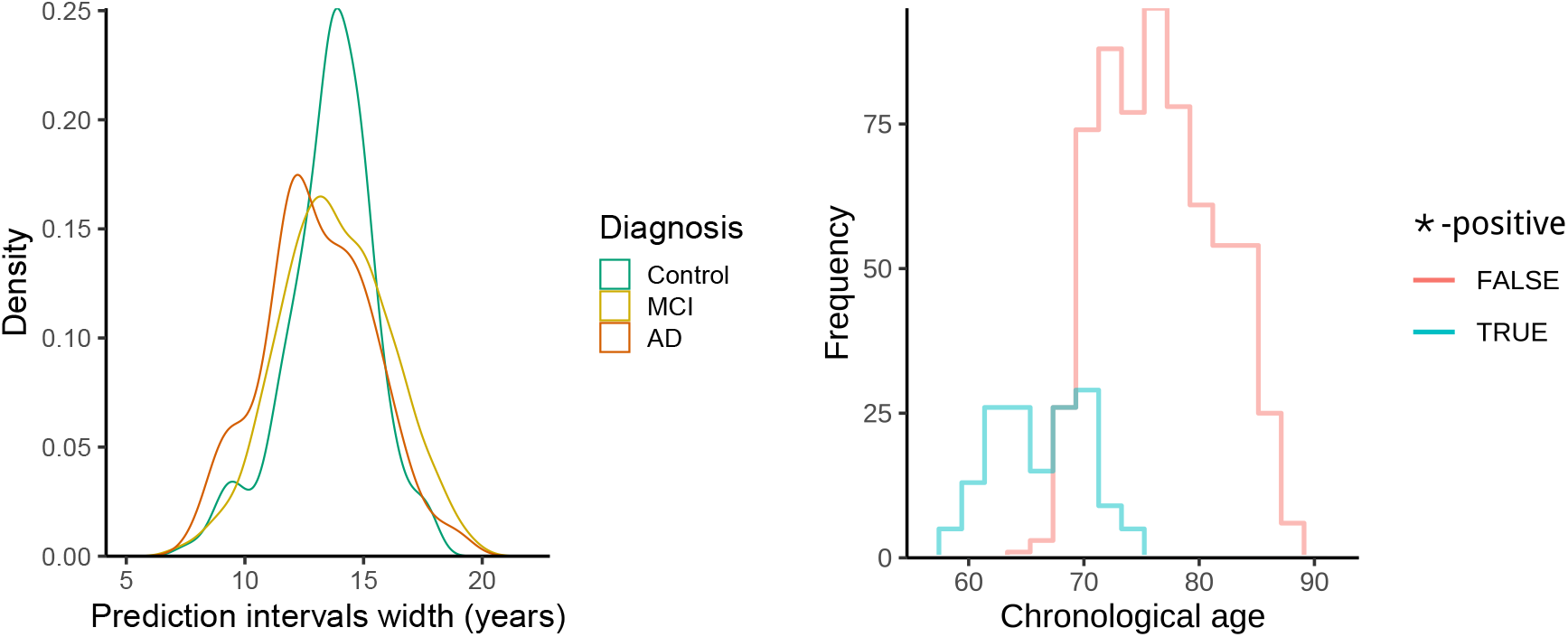
Left: distribution of the prediction interval width conditioned by diagnosis. Right: histogram of chronological age conditioned by *-positive indicator (equal to 1 if the chronological age is less than the prediction at *τ* = 0.05, 0 otherwise).

The brain maps displayed in Figure 8 are the functional coefficients obtained from the scalar-on-image quantile regression trained on the whole control dataset. They can be used to identify the regions that are responsible for the age prediction for the different quantiles. The functional coefficient for *τ* = 0.05 shows that the expansion of the lateral ventricles is the principal factor that leads to higher predicted age (Preul et al., 2006; Apostolova et al., 2012) in the lower tail of the chronological age distribution. Other areas seem to have more limited impact on the prediction. In the coefficient obtained from the median regression, the lateral ventricles still play a role in the prediction (especially the posterior part) but expansion in several other areas is correlated to higher predicted age. Among them we point out the central sulcus (perpendicular to the median longitudinal fissure that divides the two hemispheres) that separates the primary motor cortex and the primary somatosensory cortex. In addition, the frontal lobe shows negative values for the functional coefficient, meaning that expansion in this part of the brain is linked to a lower predicted age. This agrees with the literature: age-related atrophy is more pronounced in the frontal lobe (Fjell et al., 2014; Cabeza and Dennis, 2013; MacPherson and Cox, 2017) and less in the occipital lobe (Dennis and Cabeza, 2011). For *τ* = 0.95, the brain map indicates that the upper part of the cortex and the cerebellum are related to higher predicted age, while a larger left temporal lobe (in blue in the lower axial slices, it plays a role in memory and language control) is associated to younger brain age. Especially for these last two maps, asymmetry between hemispheres appears in the relationship with brain age.

**Figure 8:**
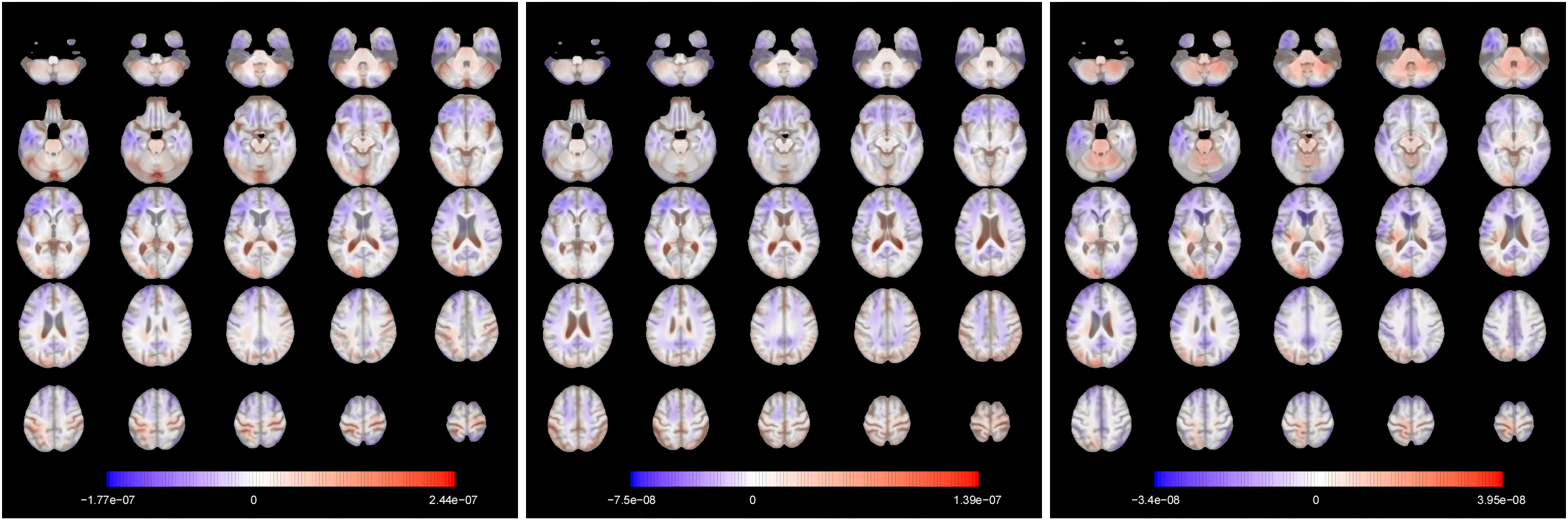
Axial slices of the functional regression coefficient for *τ* = {0.05, 0.5, 0.95} (from left to right). Slices are ordered from bottom to top. The colours are overlaid on the corresponding slice of the MDT. For a unit increase (expansion) in the observed TBM image in a red voxel, there is an increase in predicted brain age, while in a blue voxel there is a decrease.

### 4.2. Correlation with cognitive decline measures

A small number of cognitive decline measures available in ADNI has been used to evaluate the clinical utility of the predictions obtained. The list of measures reported in Table 3 includes genetic assessments (ApoE4) and various evaluations of writing and speaking skills, visual attention and task switching. The outcomes of interest in this section are both the brain-predicted age difference (*brainPAD*, difference between predicted and chronological age, as defined in Cole et al., 2017) and the binary *-positive indicator (equal to 1 if the chronological age is less than the prediction at *τ* = 0.05, 0 otherwise).

**Table 3:** Cognitive decline measures used in the analysis. The arrows indicate the change in the measures associated to an increase in dementia severity.

| Variable |  | Values |  |
| --- | --- | --- | --- |
| ApoE4 | Apolipoprotein E - Number of $\varepsilon 4$ alleles | $\{0, 1, 2\}$ | ↗ |
| ADAS11 | AD Assessment Scale - 11-item variant | $\{0, 1, \dots, 70\}$ | ↗ |
| ADAS13 | AD Assessment Scale - 13-item version | $\{0, 1, \dots, 85\}$ | ↗ |
| ADASQ4 | AD Assessment Scale - Delayed Word Recall | $\{0, 1, \dots, 10\}$ | ↗ |
| MMSE | Mini-Mental State Examination | $\{0, 1, \dots, 30\}$ | ↘ |
| DIGITSCOR | Digit Symbol Substitution Test | $\{0, 1, \dots, 83\}$ | ↘ |
| TRABSCOR | Trails B Making Test | $\{0, 1, \dots, 996\}$ | ↗ |

Figure 9 summarises the main findings in this validation analysis. A higher ApoE4 value— linked to higher risk of dementia—is also related to higher predicted age difference on average (the p-values refer to one-sided tests). In addition, for the group with the highest ApoE4, more than 75% of the individuals show higher predicted age than chronological.

**Figure 9:**
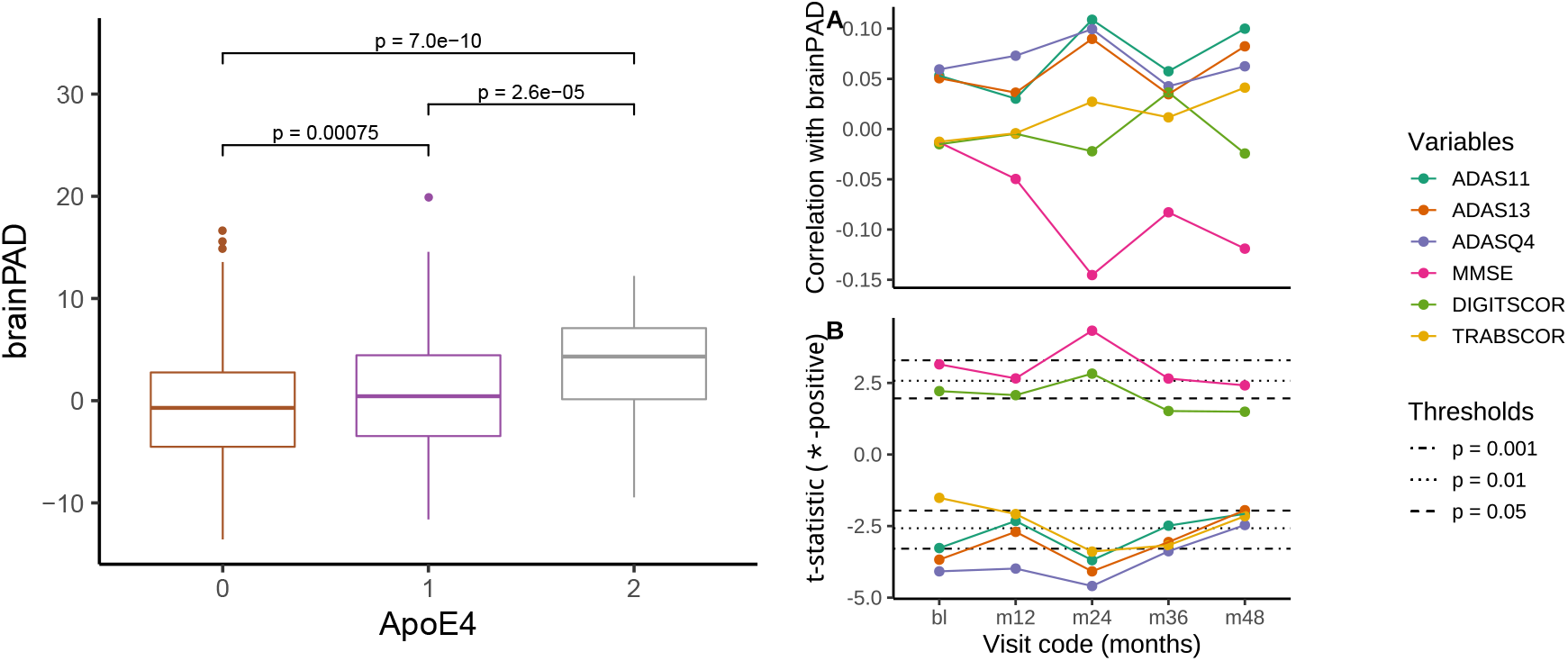
Left: association of *brainPAD* with ApoE4 value (Holm-corrected p-values) for different visits, with evidence of positive association. Right: (A) Correlation between baseline *brainPAD* and cognitive scores at different visits; (B) t-statistic for the comparisons of means of cognitive scores between *-positive group and the rest of the sample at different visits. The black lines are Student’s t quantiles which correspond to different probabilities in the tails of the distribution.

The correlation between baseline *brainPAD* and cognitive scores at different visits shows some association (uncorrected) for several measures, with ADAS measures and MMSE showing the strongest associations after 2 years. Nevertheless, no cognitive measure recorded at baseline is associated with the difference between predicted and chronological age. On the other hand, there is some evidence that the average of the cognitive measures is different between the *-positive group and the rest of the subjects across different time points. Also in this case the direction of the relationship is consistent with the numerical definition of the measures.

### 4.3. Sensitivity analysis

The prediction results are obtained under specific choices of several parameters. In order to assess how these choices might affect the results, we perform a sensitivity analysis using different values of the following parameters:

- PVE: proportion of variance explained (criterion to decide the number of fPC to be included in the quantile regression models), PVE ∈ {0.65, 0.8, 0.95};
- KS: knot spacing, KS ∈ {6, 9, 12, 15};
- nominal coverage: desired width of the prediction intervals. Values considered:

– *τ* ∈ {0.1, 0.5, 0.9} for a 80% nominal coverage,
– *τ* ∈ {0.05, 0.5, 0.95} for a 90% nominal coverage.

For each combination of values, we get the projections for each image and then fit the LASSO quantile regression. For the cases with KS = 6, the standard procedure did not work because of a failure in the Cholesky decomposition of the weight matrix *W* in Section 2.4, due to numerical tolerance issues. In these cases, the pivoted Cholesky decomposition can be applied: due to the fact that the matrix *W* is symmetric semipositive definite by construction, there is a permutation matrix *P* for which *P*^*T*^ *WP* can be factorised with an upper triangular matrix (see Higham, 2009 for an introduction).

We report as main outcomes the mean absolute error and the actual relative coverage (1 − *h*, where *h* is the ratio between observed and nominal coverage) obtained for the control subjects in Figure 10.

**Figure 10:**
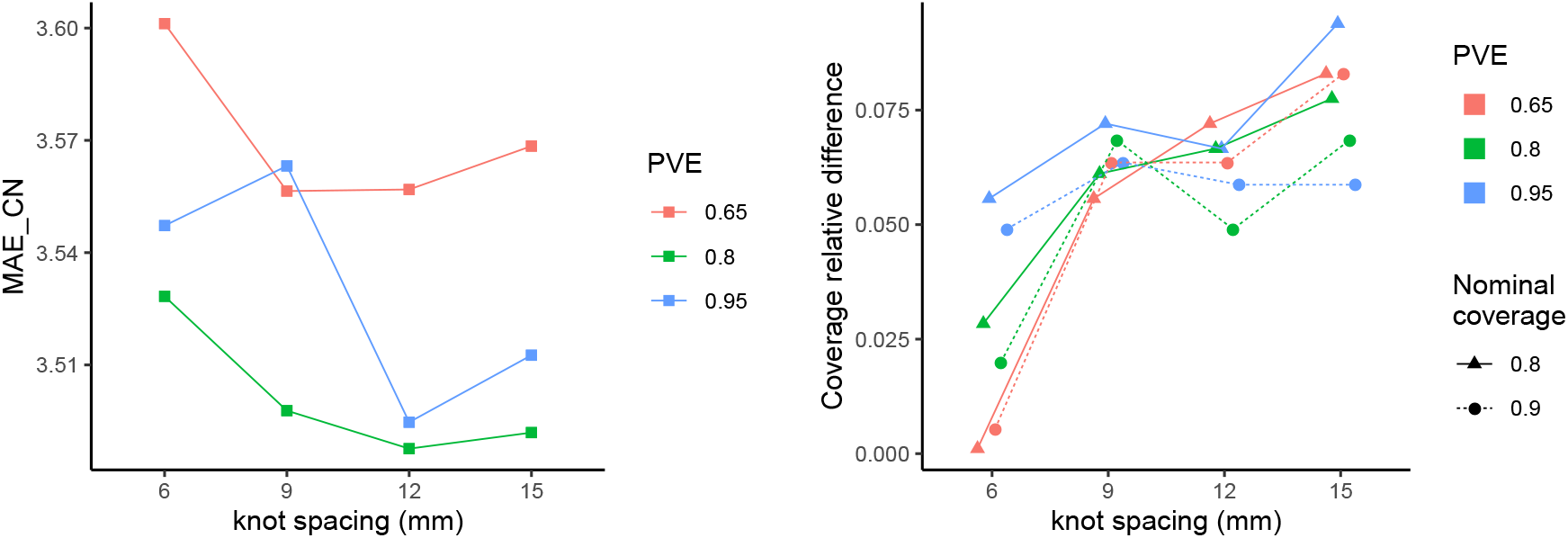
Left: mean absolute error for control subjects as function of proportion of variance explained and knot spacing. Right: Coverage relative difference of prediction intervals induced by each choice of proportion of variance explained, knot spacing and nominal coverage. Points are jittered horizontally for visualisation purposes.

The MAE refers to the predictions obtained with *τ* = 0.5, so it is not affected by the choice of nominal coverage. In general, the MAE remains rather stable across combinations of PVE and knot spacing, suggesting that our results are robust to the choices of these parameters. The lower MAE is always achieved for PVE = 0.8: this might suggest that a low PVE neglects important sources of variation while a higher one introduces too many useless variables in the models. In terms of knot spacing, 12 mm gives in almost all the cases the best results across PVE values.

Looking at the coverage for each setting of knot spacing, PVE and nominal coverage, we first observe that there are no cases in which the observed coverage is higher than the nominal level. This phenomenon of undercoverage gets more pronounced for higher knot spacing values. Except for KS = 6, when the coverage relative difference increases as the number of components in the quantile regression increases, for the other KS values no clear pattern is visible. The relative difference seems not to be influenced by the prespecified nominal coverage.

The table in the Supplementary Material section includes also a sanity check based on non-monotonic prediction intervals - those for which the predicted age at the upper *τ* level is smaller than the one at the lower level. The number of occurrences of this phenomenon is negligible in almost all the cases.

As an additional analysis, we have explored the prediction performances in terms of MAE for the control group in two models which do not use the basis expansion step, using the R packages `bigmemory` (Kane et al., 2013) and `bigstatsr` (Privé et al., 2018). The first model (M1) is a sparse linear regression with LASSO regularisation applied on the unsmoothed data (represented by 1 column per voxel in the data matrix). The second model (M2) is closer to our approach: a PCA is performed on the covariance of the matrix of unsmoothed images, then the scores corresponding to the first principal components selected (using a proportion of variance explained of at least 0.8) are plugged into a penalised quantile regression model. M2 can be interpreted as a special case of our functional approach when the distance between adjacent knots is equal to 1 mm.

The difference in computational time between our approach (M0) and the models M1 and M2 is not substantial. On one hand, the smoothing step in M0 is performed independently for each image in a parallelised setting therefore it requires only a few minutes in total. On the other hand, M1 and M2 require to load the matrix (6.4 GB in our case) in memory and run sparse linear regression or PCA and quantile regression which could take several minutes. For what concerns the prediction performances, M0 achieves lower MAE for the control group with respect to M1 (MAE = 3.63) and M2 (MAE = 3.65).

## 5. Discussion and further research

The functional data paradigm represents a useful approach to the analysis of complex data such as brain scans and offers a way to fit a global model for 3D images. In this work we have discussed the basic aspects of functional data and presented an application of quantile scalar-on-image regression (as extensions of classical quantile regression) in the field of brain age studies. Following the existing literature, we have devised an efficient workflow that takes as input a tensor-based morphometry image and returns a prediction interval. The advantages of employing the whole images as covariates are that some common preprocessing steps might be avoided (e.g. brain tissue segmentation) and there is no need to summarise information at the ROI (regions of interest) level. In addition, quantile regression gives a more detailed picture of the relationship between the covariate and the response and returns an interval with the desired coverage when the distribution of the dependent variable departs from normality. In contrast with other existing models coming from a machine learning perspective, our method outputs not only a point estimate but also a prediction interval. In addition, the model allows to investigate the functional coefficient estimated, in order to visualise the brain regions that influence most the predicted age.

Our modelling strategy introduces new features with respect to the standard prediction-oriented approaches in the literature. While other approaches focus only on maximising prediction accuracy, we emphasise the detection of individual atypical ageing: the prediction intervals give a simple and preliminary assessment of the relevance of the observed brainPAD. In other words, the same brainPAD could be indicative of potential neurodegenerative diseases for one subject, while being less linked to such disease for another subject.

The results from the analysis of ADNI data are encouraging: the point (median) prediction performances in terms of MAE and RMSE for the control subjects are comparable with the literature on the topic—even with deep learning approaches applied on bigger ADNI datasets (Varatharajah et al., 2018)—while being also more principled and interpretable. The correlation between chronological and predicted age results to be lower than the one found with other methods. The model trained on the control group highlights differences with respect to the MCI and AD groups: individuals with cognitive impairment are predicted to be older on average than their observed age, as observed in the literature (Cole et al., 2017; Franke et al., 2012).

The model proposed is an example of penalised functional regression. In this respect, some degree of regularisation can be applied at different stage of functional data analysis, starting from smoothing (Ramsay and Silverman, 2005). At the same time, the choice of the number of functional principal components to be used in regression (by using the proportion of variance explained) is itself a penalisation. On top of this we added a further penalisation, driven this time by the relationship between outcome and predictors, to account for the potential high number of covariates given the sample size (following the indication provided in Heinze et al., 2018). Our model represents a novelty in the literature as it easily accommodates this aspect into a quantile regression model with 3D functional covariates.

In addition to the bias induced by the regularisation, another potential issue related to the functional coefficient is its sensitivity to the modelling strategy used. As extensively studied in Happ et al. (2018), the smoothness induced by splines could lead to different estimates with respect to other approaches (e.g. wavelet basis expansion or random field methods). Further work can be done to confirm the contribution of each brain region to the final prediction. Nevertheless, the predictive ability - which is the first focus of our model - does not seem to be harmed by this modelling choice.

Our approach is competitive in terms of speed compared to existing methods (Franke et al., 2012; Cole, 2017). In particular, for a new image the model returns the predicted interval in approximately a minute and the training phase of the model is expected to be shorter and less computationally intensive than training a neural network, especially because the basis expansion step runs in parallel for each image.

The modelling approach illustrated in this paper can be extended in multiple ways, from both theoretical and practical perspectives. For what concerns the key points of the workflow, in this paper we have chosen to project the images (and the functional coefficients) using B-spline basis functions and sketched a possible strategy to select knot spacing. We have shown that some degree of smoothing produces slightly better predictions with respect to no smoothing at all with negligible computational cost. The benefit of this approach could more easily appreciated when the number of images is much larger, in which case loading the whole unsmoothed data into memory can be unfeasible.

The quantile regression approach is a technically easy-to-implement strategy to build prediction intervals without assuming normality. Since we consider only the best fit for each of the regression models, it could be of interest to study how the uncertainty about the coefficients and the models could play a role in the calculation of individual prediction intervals. The observed coverage in the control group could also depend on the bias/variance trade-off introduced by the cross-validation procedure (and in particular on the type of penalty and the number of folds chosen). Further simulation study can be done to assess the extent of this relationship.

In addition, further extensions of quantile regression could be considered. Additive terms might be introduced in order to explore nonlinear effects of the imaging covariate. Moreover, quantile boosting (Mayr et al., 2012) could provide better prediction intervals by reducing the bias due to the estimation at extreme quantiles. This approach has a higher computational cost but keeps the advantage of interpretability, which is no longer available with other approaches such as quantile regression forests described in Meinshausen, 2006. A potential issue for the current formulation of our approach is the phenomenon of *quantile crossing*, that occurs when the predicted quantiles are not monotonically increasing in *τ* as the conditional quantile function is by construction. Although in 90% prediction intervals the problem arises rarely (in our application it has been reported for only 1 case out of 796), still this could introduce some bias. Monotonicity can be forced after the estimation by using rearrangement or isotonic regression (see e.g. Kato, 2012; Chernozhukov et al., 2010). An alternative modelling strategy for quantile regression that ensures monotonicity of the function is provided in Chen and Müller (2012): the quantile function is obtained indirectly by first estimating the entire CDF of the response variable and then inverting it to recover the quantile function at the level of interest. The key idea is to use a generalised functional linear model to model the conditional distribution of *Y* |*X* as conditional expected values of indicator functions. This “indirect” model is claimed to provide better estimation of the quantile function with respect to the classical quantile regression at extreme quantile levels for non-gaussian response variables (Chen and Müller, 2012), although the flexibility induced by considering different predictors at different quantile levels is lost. In addition, generalised additive models for location, scale and shape (GAMLSS, Rigby and Stasinopoulos (2005)) can also provide a detail picture of the conditional distribution of the outcome of interest. In GAMLSS the parameters of the distribution (not only the location, as in GLM) can be written as (smooth) functions of the covariates. GAMLSS can handle functional covariates (Brockhaus et al., 2018) and ensures monotonocity of the quantile predictions, but the family of the conditional distribution of the outcome must be specified in advance.

From the application point of view, it is currently very difficult to provide a sensible comparison between different models. This is due to the large range of possible approaches (from multivariate statistics to deep learning) applied to a plethora of datasets with different sizes, age ranges and imaging modalities (T1-weighted MRI to PET or FMRI). Cole et al. (2019) uses a MAE weighted by the age range in the training set as a measure of comparison. That approach might be too simplistic, as a 1-year absolute error for a 6-year child should probably be weighted more than the same error for a 70-year old individual. A more adaptive measure should be devised, or alternatively there should be an incentive towards the use of a specific dataset as a benchmark. Big databases such as UK Biobank (Sudlow et al., 2015) seem the right testing ground for all the methods available in the literature. Our model could be applied on different imaging modalities, for example voxel-based morphometry, in order to specify potential differences in the effects due to white and gray matter. Coming to more specific modelling-related issues, as observed from the plots concerning the prediction intervals, a non negligible correlation is noticed between chronological age and the brain age differences (predicted minus chronological, called *brainPAD* in Cole et al., 2017, *brainAGE* - brain age gap estimate - in Franke and Gaser, 2019 or *δ* in Smith et al., 2019). This undesirable effect arises from the simple fact that by construction the residuals (which become the objects of interest when we want to explore the relationship with other variables such as disease conversion) in a regression model are uncorrelated with respect to the predicted values, but not with the observed ones. Similar issues are also reported in the deep learning approaches to brain age prediction (Cole et al., 2017; Varatharajah et al., 2018). The work by Smith et al. (2019) identifies potential reasons for this phenomenon and proposes some solutions. Among others, a viewpoint that is conceptually grounded and at the same time can be embedded in our model could be rephrasing the whole problem in terms of a errors-in-variables framework. In particular, this accounts for the imaging covariate (consistently with the functional data perspective) or its scores representation being measured with some errors. At the same time, the response itself (chronological age) can be considered as a noisy proxy for biological brain age (for which it is difficult or even impossible to define a reference measure).

Another aspect left for future research is to extend the analysis of the clinical utility of the prediction intervals obtained with our workflow by using a larger battery of cognitive measures. The first basic measures selected in this work show interesting and sensible results, especially for the correlation with the *-positive binary variable. A desired feature of this indicator in a prognostic context should be its correlation with conversion to dementia, in order to provide a sensible way to early detect neurodegenerative diseases. Furthermore, a similarly defined “*-negative indicator” could be also explored in the same way in order to show potential aspects of a healthy aging process.

In addition, introducing other covariates in the model (such as sex, years of education or physical activity measures) is rather straightforward and it could improve the detection of discrepancies from normative ageing. On the other hand, these covariates might potentially introduce confounding effects: the variability due to non-imaging information could be already captured by one or more functional principal components. Our approach can be also easily incorporated in a longitudinal model where brain age trajectories could provide evidence of stable or accelerated brain ageing.

## Acknowledgements and funding sources

MP is funded by the EPSRC and MRC Centre for Doctoral Training in Next Generation Statistical Science: The Oxford-Warwick Statistics Programme (EP/L016710/1). TEN is supported by the Wellcome Trust, 100309/Z/12/Z. We would like to thank David Firth, Xavier Didelot, Ioannis Kosmidis and the anonymous reviewers for their insightful comments about the work. We also thank Paul Thompson and Xue Hua for the TBM data. Data collection and sharing for this project was funded by the Alzheimer’s Disease Neuroimaging Initiative (ADNI) (National Institutes of Health Grant U01 AG024904) and DOD ADNI (Department of Defense award number W81XWH-12-2-0012). ADNI is funded by the National Institute on Aging, the National Institute of Biomedical Imaging and Bioengineering, and through generous contributions from the following: AbbVie, Alzheimer’s Association; Alzheimer’s Drug Discovery Foundation; Araclon Biotech; BioClinica, Inc.; Biogen; Bristol-Myers Squibb Company; CereSpir, Inc.; Cogstate;Eisai Inc.; Elan Pharmaceuticals, Inc.; Eli Lilly and Company; EuroImmun; F. Hoffmann-La Roche Ltd and its affiliated company Genentech, Inc.; Fujirebio; GE Healthcare; IXICO Ltd.; Janssen Alzheimer Immunotherapy Re-search & Development, LLC.; Johnson & Johnson Pharmaceutical Research & Development LLC.; Lumosity; Lundbeck; Merck & Co., Inc.; Meso Scale Diagnostics, LLC.; NeuroRx Research; Neurotrack Technologies;Novartis Pharmaceuticals Corporation; Pfizer Inc.; Piramal Imaging; Servier; Takeda Pharmaceutical Company; and Transition Therapeutics. The Canadian Institutes of Health Research is providing funds to support ADNI clinical sites in Canada. Private sector contributions are facilitated by the Foundation for the National Institutes of Health (www.fnih.org). The grantee organization is the Northern California Institute for Research and Education, and the study is coordinated by the Alzheimer’s Therapeutic Research Institute at the University of Southern California. ADNI data are disseminated by the Laboratory for Neuro Imaging at the University of Southern California.

## Footnotes

2 the word “functional” in this case is used in a mathematical sense and is not related to functional MRI.

3 From Mosteller and Tukey (1977): ‘Just as the mean gives an incomplete picture of a single distribution, so the regression curve gives a correspondingly incomplete picture for a set of distributions.’

