## supplementary for "Quantifying uncertainty in brain-predicted age using scalar-on-image quantile regression" (from another version of this preprint: version 1, 2019-11-26)

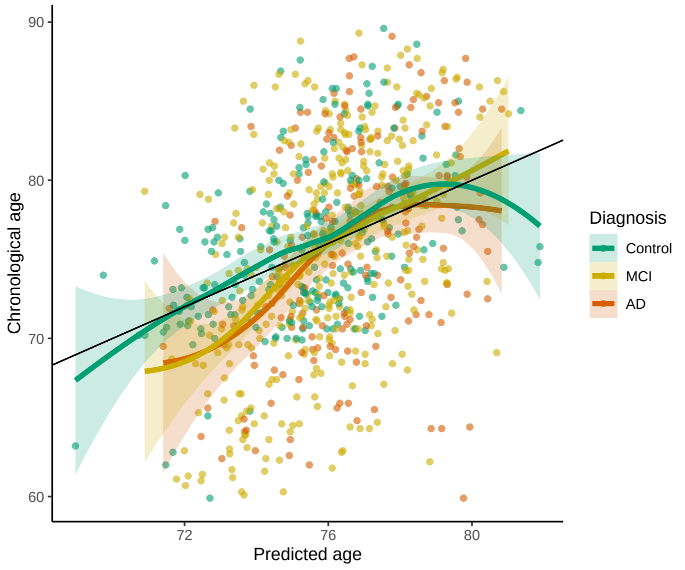


Figure S.1: **Plot of the observed vs. predicted response.** The black line is the line of identity $y = x$. The coloured lines are local regression lines obtained with loess (locally estimated scatterplot smoothing) with span = 0.75 and 95% confidence bands.

| nominal coverage | pve | KS | MAE_CN | MAE_MCI | MAE_AD | Cover_CN | Cover_MCI | Cover_AD | non monotone intervals | # basis | density percentage | #fPC |
|---|---|---|---|---|---|---|---|---|---|---|---|---|
| 0.8 | 0.65 | 6 | 3.601 | 5.086 | 5.228 | 0.799 | 0.636 | 0.622 | 0 | 14591 | 0.156 | 18 |
| 0.9 | 0.65 | 6 | 3.601 | 5.086 | 5.228 | 0.895 | 0.687 | 0.694 | 3 | 14591 | 0.156 | 18 |
| 0.8 | 0.80 | 6 | 3.528 | 5.065 | 5.148 | 0.777 | 0.618 | 0.583 | 0 | 14591 | 0.156 | 28 |
| 0.9 | 0.80 | 6 | 3.528 | 5.065 | 5.148 | 0.882 | 0.685 | 0.672 | 1 | 14591 | 0.156 | 28 |
| 0.8 | 0.95 | 6 | 3.547 | 5.003 | 5.315 | 0.755 | 0.574 | 0.561 | 2 | 14591 | 0.156 | 55 |
| 0.9 | 0.95 | 6 | 3.547 | 5.003 | 5.315 | 0.856 | 0.687 | 0.633 | 1 | 14591 | 0.156 | 55 |
| 0.8 | 0.65 | 9 | 3.556 | 5.016 | 5.167 | 0.755 | 0.561 | 0.539 | 1 | 5294 | 0.455 | 27 |
| 0.9 | 0.65 | 9 | 3.556 | 5.016 | 5.167 | 0.843 | 0.705 | 0.722 | 0 | 5294 | 0.455 | 27 |
| 0.8 | 0.80 | 9 | 3.498 | 5.001 | 5.156 | 0.751 | 0.589 | 0.561 | 0 | 5294 | 0.455 | 59 |
| 0.9 | 0.80 | 9 | 3.498 | 5.001 | 5.156 | 0.838 | 0.682 | 0.661 | 1 | 5294 | 0.455 | 59 |
| 0.8 | 0.95 | 9 | 3.563 | 5.001 | 5.156 | 0.742 | 0.592 | 0.561 | 2 | 5294 | 0.455 | 143 |
| 0.9 | 0.95 | 9 | 3.563 | 5.001 | 5.156 | 0.843 | 0.682 | 0.661 | 1 | 5294 | 0.455 | 143 |
| 0.8 | 0.65 | 12 | 3.557 | 5.016 | 5.166 | 0.742 | 0.579 | 0.578 | 1 | 2694 | 0.921 | 26 |
| 0.9 | 0.65 | 12 | 3.557 | 5.016 | 5.166 | 0.843 | 0.705 | 0.722 | 0 | 2694 | 0.921 | 26 |
| 0.8 | 0.80 | 12 | 3.488 | 4.994 | 5.156 | 0.747 | 0.592 | 0.561 | 0 | 2694 | 0.921 | 57 |
| 0.9 | 0.80 | 12 | 3.488 | 4.994 | 5.156 | 0.856 | 0.680 | 0.644 | 1 | 2694 | 0.921 | 57 |
| 0.8 | 0.95 | 12 | 3.495 | 5.001 | 5.158 | 0.747 | 0.589 | 0.567 | 0 | 2694 | 0.921 | 138 |
| 0.9 | 0.95 | 12 | 3.495 | 5.001 | 5.158 | 0.847 | 0.674 | 0.661 | 1 | 2694 | 0.921 | 138 |
| 0.8 | 0.65 | 15 | 3.568 | 5.022 | 5.170 | 0.734 | 0.581 | 0.567 | 1 | 1617 | 1.561 | 24 |
| 0.9 | 0.65 | 15 | 3.568 | 5.022 | 5.170 | 0.825 | 0.630 | 0.622 | 11 | 1617 | 1.561 | 24 |
| 0.8 | 0.80 | 15 | 3.492 | 5.007 | 5.160 | 0.738 | 0.589 | 0.567 | 0 | 1617 | 1.561 | 54 |
| 0.9 | 0.80 | 15 | 3.492 | 5.007 | 5.160 | 0.838 | 0.687 | 0.656 | 1 | 1617 | 1.561 | 54 |
| 0.8 | 0.95 | 15 | 3.513 | 5.017 | 5.173 | 0.725 | 0.587 | 0.567 | 0 | 1617 | 1.561 | 132 |
| 0.9 | 0.95 | 15 | 3.513 | 5.017 | 5.173 | 0.847 | 0.687 | 0.656 | 1 | 1617 | 1.561 | 132 |

Table S.1: Results from the sensitivity analysis. PVE: proportion of variance explained (criterion to decide the number of fPC to be included in the quantile regression models). KS: knot spacing. Nominal coverage: desired width of the prediction intervals (if 0.8, $\tau = \{0.1, 0.5, 0.9\}$, if 0.9, $\tau = \{0.05, 0.5, 0.95\}$). Non monotone intervals: number of intervals for which 0.95 prediction ¡ 0.05 prediction. Density percentage: number of nonzero values in the basis design matrix (with dimensions $\#voxels \times \#basis$)
